# Assessing Attentiveness and Cognitive Engagement across Tasks using Video-based Action Understanding in Non-Human Primates

**DOI:** 10.1101/2025.05.31.657183

**Authors:** Sin-Man Cheung, Adam Neumann, Thilo Womelsdorf

## Abstract

**Background:** Distractibility and attentiveness are cognitive states that are expressed through observable behavior. The effective use of behavior observed in videos to diagnose periods of distractibility and attentiveness is still not well understood. Video-based tools for classifying cognitive states from behavior have high potential to serve as versatile diagnostic indicators of maladaptive cognition.

**New method:** We describe an analysis pipeline that classifies cognitive states using a 2-camera set-up for video-based estimation of attentiveness and screen engagement in nonhuman primates performing cognitive tasks. The procedure reconstructs 3D poses from 2D labeled DeepLabCut videos, reconstructs the head/yaw orientation relative to a task screen, and arm/hand/wrist engagements with task objects, to segment behavior into an attentiveness and engagement score.

**Results:** Performance of different cognitive tasks were robustly classified from video within a few frames, reaching >90% decoding accuracy with ≤3min time segments. The analysis procedure allows setting subject-specific thresholds for segmenting subject specific movements for a time-resolved scoring of attentiveness and screen engagement.

**Comparison with existing methods:** Current methods also extract poses and segment action units; however, they haven’t been combined into a framework that enables subject-adjusted thresholding for specific task contexts. This integration is needed for inferring cognitive state variables and differentiating performance across various tasks.

**Conclusion:** The proposed method integrates video segmentation, scoring of attentiveness and screen engagement, and classification of task performance at high temporal resolution. This integrated framework provides a tool for assessing attention functions from video.

## 1. Introduction

A goal of behavioral neuroscience is to distinguish cognitive states from behavior. One approach to achieve this goal is to use video to detect and understand behavioral patterns that are diagnostic for maladaptive cognition. Such an action understanding from video requires analysis procedures for extracting behavioral patterns from video while subjects engage in cognitive demanding tasks (Vogg et al., 2025). Recent examples have showcased the potential clinical value of evaluating of maladaptive cognitive states with video based analysis, ranging from classifying behavioral grooming abnormities in animals with fragile X syndrome (Marshall et al., 2021), to the video-based detection of drug induced changes in behavioral speed in mice (Wiltschko et al., 2020). These advances have been enabled by video-based machine learning tools that allow segmenting natural behavior into action units such as DeepLabCut (Mathis et al., 2018), OpenMonkeyStudio (Bala et al., 2020; Bain et al., 2021), MacaquePose (Labuguen et al., 2020), LightPose (Biderman et al., 2024), Deepercut (Insafutdinov et al., 2016), SLEAP (Pereira et al., 2022), or Convolutional ‘Deep’ Pose Machine (Wei et al., 2016). While these tools have optimized the 3D reconstruction of behavioral patterns it has remained rare to relate free-ranging behaviors of subjects to specific cognitive factors such as the degree of attentiveness or the sustained attentional engagement with differently demanding tasks (Marshall et al., 2022). Consequently, it has remained a challenge to individualize video-based behavioral analysis that quantifies cognitive factors in nonhuman primates (Vogg et al., 2025).

Here, we set out to establish an integrated analysis procedure that combines video-based machine learning segmentation of head, shoulder, arm, wrist, and hand of rhesus monkeys with a subject-specific evaluation of attentiveness and screen engagement during performance of multiple cognitive tasks. Attentiveness describes the vigilance and sustained attention subjects show during goal-directed behavior. Quantifying attentiveness has highest clinical significance because it is affected not only in attention-deficit disorders (Fuermaier et al., 2022), but across all major neuropsychiatric disorders (Millan et al., 2012; Grant and Chamberlain, 2023). Evaluating attentiveness from video requires measuring the consistency of head and gaze orientation towards task-relevant stimuli. Previous studies have inferred gaze and head orientation from video in marmosets using ear and face landmarks (Xing et al., 2024), head-mounted gaze tracking devices (Kano and Tomonaga, 2013; Singh et al., 2025), or by using boxes that restrict the field of view (Ryan et al., 2019). These approaches have been used to characterize the rich gaze behavior of animals with high-resolution foveal (Singh et al., 2025), but this sensorimotor variables of head and gaze orientation have not been used to infer cognitive variables. The potential to infer more cognitive variables has been documented by the rodent field where gaze and head behavioral patterns have been linked to decision making (Redish, 2016), reward-based learning (Ottenheimer et al., 2020), or multi-sensory integration (Keshavarzi et al., 2022). Here, we aimed at using head, arm and body orientation to quantify an attentiveness score and determine how it varies over time when NHPs engage with multiple tasks with varying cognitive demands.

A second aim of this study is to distinguish cognitive engagement in different tasks. Engagement corresponds to the degree of interacting with task-relevant stimuli, which requires wrist and hand movements towards a screen. Previous studies in primates succeeded to segment such hand and digit movements during the manipulation of reward boxes (Voloh et al., 2023), during nut cracking and buttress drumming (Bain et al., 2021), feeding behaviors (Vogg et al., 2025), precision grasping (North et al., 2021; Li et al., 2023; Liu et al., 2025), or as part of standard sets of daily activity together with sitting, walking or climbing behaviors (Brookes et al., 2024). While these studies use hand or digit reconstruction, they have not yet been applied to touchscreen engagement during different cognitive tasks.

Here, we address the open questions on how task engagement and attentiveness can be tracked from video in individual monkeys in order to distinguish their performance of different cognitive tasks. We report of an integrated analysis pipeline that uses state-of-the-art pose estimation together with custom reconstruction of head, arm and hand orientation towards a touchscreen in order to derive a frame-by-frame estimate of attentiveness and task engagement. We show that these cognitive scores allow distinguishing which of three different cognitive tasks are performed by NHPs in a touchscreen Kiosk setting in their home cage.

## 2. Methods and Analysis

### 2.1. Experimental set-up

All animal and experimental procedures complied with the National Institutes of Health Guide for the Care and Use of Laboratory Animals and the Society for Neuroscience Guidelines and Policies and were approved with the approval number M1700198 by the institutional review board Vanderbilt University Institutional Animal Care and Use Committee (IACUC). The experiment involved four nonhuman primates (NHPs, rhesus monkeys, 11.5-12.5 kg) who performed cognitive tasks using a touchscreen kiosk station that was mounted to the home cage. The kiosk station, described in detail in (Womelsdorf et al., 2021), was equipped with two video cameras and a reward pump connected to a stainless steel sipper tube placed ∼30’’ in front of the touchscreen. The cognitive tasks were controlled by the *Multi-task Suite for Experiments* (<u>M-USE</u>), an open source unity-based behavioral task platform designed to control the timing of visual stimuli, registering responses of subjects and deliver fluid reward (Watson et al., 2023).

### 2.2. Cognitive task paradigms

In ∼96 min long daily experimental sessions, subjects engaged with five separate cognitive tasks in a fixed temporal order: (1) Maze-Learning task part 1 (MZ1, ∼18 min), (2) delayed match-to-sample Working Memory task part 1 (WM1, ∼20 min), (3) Effort Control task (EC, ∼20 min), (4) delayed match-to-sample Working part 2 (WM2, 20 min), and (5) Maze-Learning task part 2 (MZ2, ∼18 min). The tasks are described in detail in (Watson et al., 2023). In brief, the first task, the Maze-Learning task, presented a 6x6 grid of tiles in which subjects were required to find an invisible path with 3-4 turns and a length of 12-14 tiles between a start tile to an end tile. The start and end tiles were colored yellow and blue, respectively, while all other tiles were grey and only transiently changed for 0.3 s to green or black when they were touched and either part of the correct path (green) or else were not on the path (black). To identify the correct sequence (path) of tiles that connect start and end tile subjects followed two rules: (1) choose among the next right, left, up, or down tiles relative to the last correctly identified tile of the path, and (2) after each wrong choice, re-touch the last correct tile of the path. Subjects learned by trial-and-error which tiles were part of the hidden path. They received immediate visual feedback and a progress step of a slider bar for correct touches (transient green coloring of the tile, forward slider step) and for incorrect touches (transient black coloring of a tile, backward slider step). Subjects received fluid reward when they completed 5 correct tiles and when the path was completed. To complete a path, subject had a maximum of 120 s but always completed them before the maximal allowed time. Each maze (i.e. each path) was repeated eight times to track how subjects learned the path by reducing erroneous choices (Watson et al., 2023).

The second task was a delayed match-to-sample working memory (WM) and involved 120 separate trials and has been validated in (Wen et al., 2025). In each trial a sample stimulus was shown for 0.5 s followed by a variable delay of 0.5-5 s void of visual stimuli, and a 0.5 s display of three objects and randomly assigned positions, one of which was the probe object. When subjects correctly chose the probe object that matched the sample object, they received visual feedback (a yellow halo around the object) and a visual token (a green coin symbol) that was added to a token bar on top of the screen). Incorrect choices resulted in a cyan colored halo and no change of tokens. Each trial used a novel set of multidimensional 3D-rendered so-called Quaddle objects that had unique body shapes, body patterns, arm types and colors (Watson et al., 2019).

The third task was an effort control task (EC) involving 60 separate trials. Each trial had an early decision phase that presented one ballon outline on the left, and on outline on the right side of the screen. The number of outlines within each balloon differed and indicated how often the balloon outline needed to be touched to virtually inflate the balloon to its full extent. The number of touches correspond to the effort the subjects must exert to inflate the balloon. On top of each ballon there were different amounts of gold-colored token coins visible that corresponded to the amount of water drops subjects would receive when the ballon below the tokens was chosen and inflated to its full extend. Once subjects chose one of the ballons by touching in its inside, that balloon was moved to the center of the screen and the other nonchosen option was removed from the screen. The subjects had to repeatedly touch inside the ballon to inflate and virtually pop the ballon to receive fluid reward. The task assesses how much reward (number of tokens) is needed to motivate subjects to choose the balloon that required more effort (i.e. more touches) to fully inflate.

Following the effort control task, the fourth task in each experimental session was the working memory task with another 120 trials. The fifth task of the session was the same Maze-Learning task as at the beginning of these session with three novel and two repeated mazes. Subjects performed all five tasks in a fixed order in every session. Individual session lasted ∼96 min. The behavioral performance was stable across all sessions analyzed in this study.

### 2.3. Video monitoring set-up

Subjects were video monitored with two e3Vision cameras (*White Matter LLC*, n.d.). The resolution was set to 1600 x 1200, recording at a rate of 30 frames per second. Two cameras were set up in stereo configuration at the top of the kiosk, with one on the left and one on the right. The lens attached to the cameras was the SL183 Lens (*Theia’s SL183/ML183 LOT Lenses - Theia Technologies*, n.d.), developed by Theia Tech. Camera synchronization signals were provided through external TTL pulses from the M-USE Unity control software to synchronize the camera with the monitor frames during cognitive task performance. The cameras were calibrated to find their intrinsic and extrinsic parameters. The calibration was done with MATLAB’s Stereo Camera Calibrator Toolbox (*Stereo Camera Calibrator*, n.d.). Fifteen checkerboard images with unique positions were selected manually and inputted into the application. The application returned a stereo parameter with the intrinsic and extrinsic parameters, ready to be triangulated.

### 2.4. 3D reconstruction of physical environment

Subjects engaged with the cognitive tasks in front of a touchscreen and a reward sipper tube. This task engagement environment was reconstructed in 3D to validate the intrinsic and extrinsic parameters of the calibration step and to provide landmarks and boundaries for analysis (**Suppl. Figure S1**). Code for reconstruction and analysis is provided freely online (*see* Appendix). Pixel coordinates of three points on the reward tube (tip, curve point, and base of the reward sipper tube) were manually extracted from the left and right camera footage and triangulated to obtain the coordinate points of the reward sipper tube. The pixel coordinates of the corners of the flat surface under the reward sipper tube were also extracted. These pixel coordinates were triangulated, and the distances between each point were calculated to compare to real-world measurements. The differences between the calculated and the measured points were minimal, confirming the validity of the parameters. The touchscreen dimensions were measured and plotted in relation to the reward sipper tube and base of the Kiosk surface (**Suppl. Figure S1**). The kiosk environment and camera angles did not vary over the course of the experiment.

### 2.5. Pose estimation and extraction of 3D coordinates of subjects

Step 1 of the data analysis was the pose estimation (**Figure 1A**). Pose estimation of NHP body parts such as the nose, eyes, elbows, wrists, and fingers was performed using DeepLabCut, a deep learning tool that enables markerless tracking of user-defined features from video data (Mathis et al., 2018). The version of DLC employed in the analysis was 2.3.10. A comprehensive DLC training pipeline was created for robust analysis (**Figure 1B**). Two DLC neural networks were created for pose estimation from the left camera and the right camera to reconstruct the NHP in a 3D space. Specifically, around 180 frames from 4 to 5 unique videos of each of 4 NHPs were labeled for both the left and the right camera, resulting in 360 frames per camera. 11 body parts were labeled, including the nose, left eye, right eye, left elbow, right elbow, left wrist, right wrist, left center knuckle, right center knuckle, left pointer finger, and right pointer finger. 95% of the frames were split for training. A ResNet-50-based neural network was trained for 30,000 iterations with Google Colab’s L4 GPU. To improve network accuracy, 50 outlier frames were extracted, relabeled, and fed back into the training. Over 3 iteration cycles, the right camera had a training error of 3.08 pixels and a test error of 9.72 pixels with pcutoff of 0.6, while the left camera had a train error of 3.9 pixels and a test error of 8.32 pixels with pcutoff of 0.6. Thereby, the errors are given by the average distances between the labels by DLC and the scorer. An example frame of labeled vs DLC predicted is shown in **Supplementary Figure S2A-B.**

**Figure 1.**
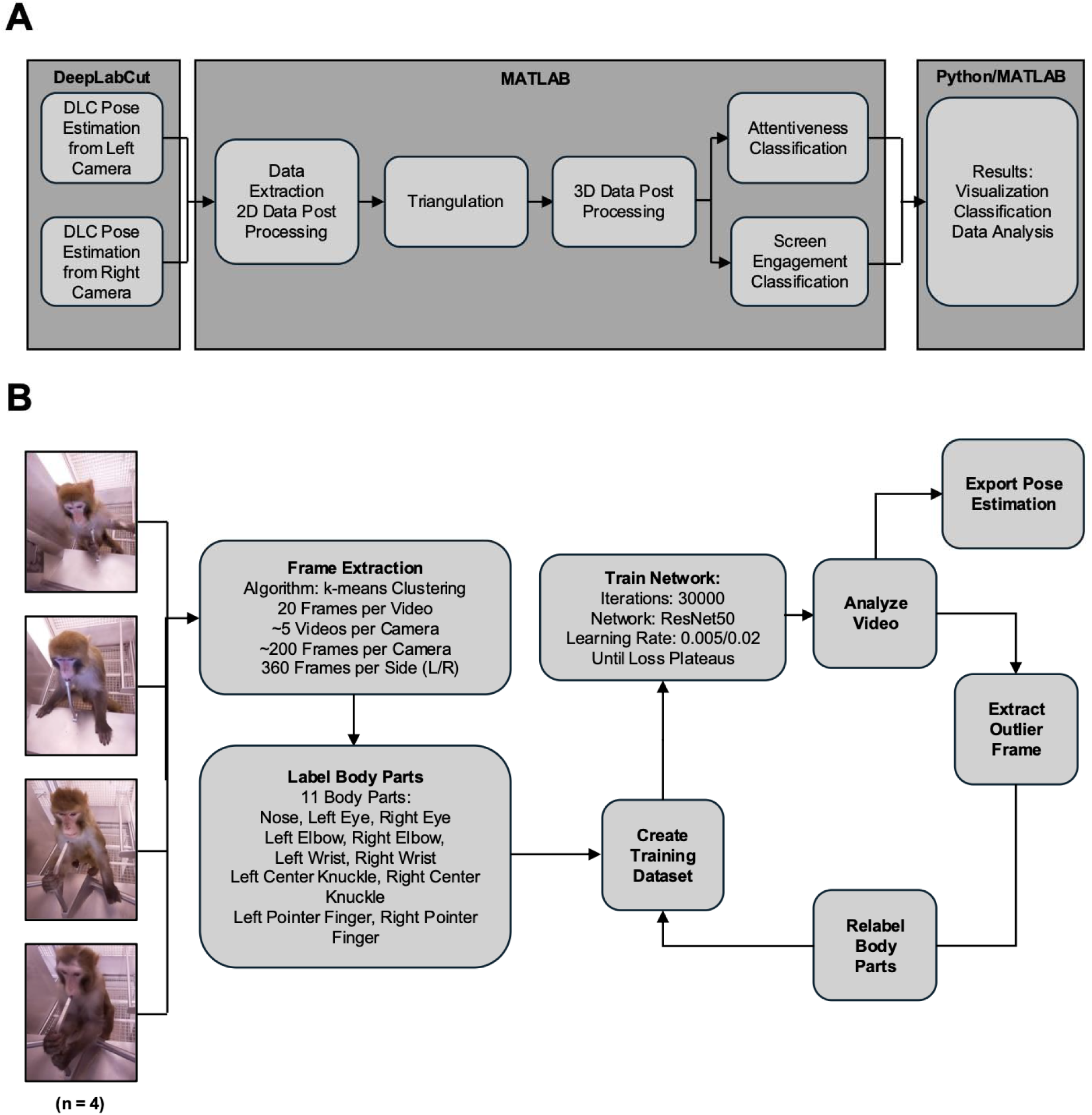
Procedural Pipelines. (A) Workflow for 3D pose estimation and classification using DeepLabCut and MATLAB/Python. Pose estimation from left and right cameras is processed via DeepLabCut, followed by 2D data extraction, triangulation, and 3D data post-processing in MATLAB. Attentiveness and screen engagement are classified, with results visualized and analyzed using Python/MATLAB. (B) Pipeline for pose estimation of rhesus macaques using DeepLabCut. Video frames (n=4 subjects) are extracted using k-means clustering (20 frames/video, 5 videos/camera, 360 frames/side). Eleven body parts are labeled, and a training dataset is created. The ResNet50 network is trained (30,000 iterations, learning rate 0.005/0.002) until loss plateaus. Outlier frames are extracted, relabeled, and used to export pose estimation for video analysis.

Step 2 of the analysis pipeline was the 3D reconstruction of the NHP subjects, illustrated in **Supplementary Figure S2**. The exported CSV data for each camera first underwent 2D post-processing in MATLAB. Low confidence data (likelihood < 0.4) was filtered out and outliers (z-score >2.5) were removed for each body part. The filtered data was then interpolated with shape-preserving piecewise cubic interpolation (pchip). The 2D data then was filtered by velocity (maximum 40 pixels/frame) and interpolated again with pchip. The 2D post-processing eliminates sudden jumps in coordinates, dropouts, and inconsistent estimations, enhancing the robustness of the data. However, the resulting data may still have erroneous segments due to the limit of interpolation. Therefore, 3D post-processing was applied. The data from the left and right cameras were triangulated with parameters obtained in the camera calibration step, transforming the data from 2D-pixel data into 3D coordinate data. The 3D data were first filtered by boundary, where points outside of the set boundary and trajectory to those points are removed and interpolated. The 3D data then was filtered by velocity (maximum 40 millimeters/frame) and interpolated with pchip. The resulting 3D coordinate data and the 3D reconstructed subject is illustrated in **Supplementary Figure S2C-F**.

### 2.6. Classification of attentiveness

Step’s 3 and 4 of the analysis-pipeline establishes the classification of attentiveness and screen engagement from the 3D frame coordinates. The classification of attentiveness and screen engagement was based on NHPs’ head orientation and movement. The primary features used for this classification include the pitch, yaw, and roll angles derived from the pose estimation of the NHP’s face (Bala et al., 2020). Specifically, the plane formed by the eyes and nose is utilized to measure the relative position of the head to the screen. These angles were computed frame-by-frame to capture dynamic changes in the NHP’s orientation. For attentiveness classification, the pitch, up and down head movement, and the yaw, left and right head movement, are the key indicators. When the pitch and yaw angles remain within specific thresholds, the NHP is classified as attentive, assuming it is looking directly at the screen. Conversely, when these angles exceed the set thresholds, the NHP is classified as inattentive, which corresponds to the subject looking away from the screen. The thresholds were adjusted individually for each NHP based on their unique behavior patterns observed during the study. This individualized approach accounts for variability in head movements and improves the accuracy of the attentiveness classification. **Figure 2A** illustrates an example analysis of an NHP’s attentiveness over a 2-minute video segment. In this analysis, the yaw angle (indicated by the black line) serves as the primary feature for determining attentiveness, with the red horizontal lines representing the left and right yaw thresholds. These thresholds served as the cutoff points to distinguish between attentive and inattentive states. In the figure, segments where the yaw angle (black line) remain within the threshold bounds indicate that the NHP is oriented toward the screen, suggesting attentiveness. These periods are marked as green shaded areas. When the yaw angle exceeds the set thresholds, the NHP is classified as inattentive. The image sequence provides snapshots from specific frames corresponding to key moments in the yaw angle trace (**Figure 2A**). For example, frames 640, 650, and 660 show the NHP transitioning from looking at the screen to looking away. In contrast, frames 740, 750, and 760 display the NHP not attentive. Similarly, frames 2680 to 2700 capture the NHP’s head turning sharply to the left, which is reflected in the yaw trace crossing the left threshold and transitioning to the grey bar.

**Figure 2.**
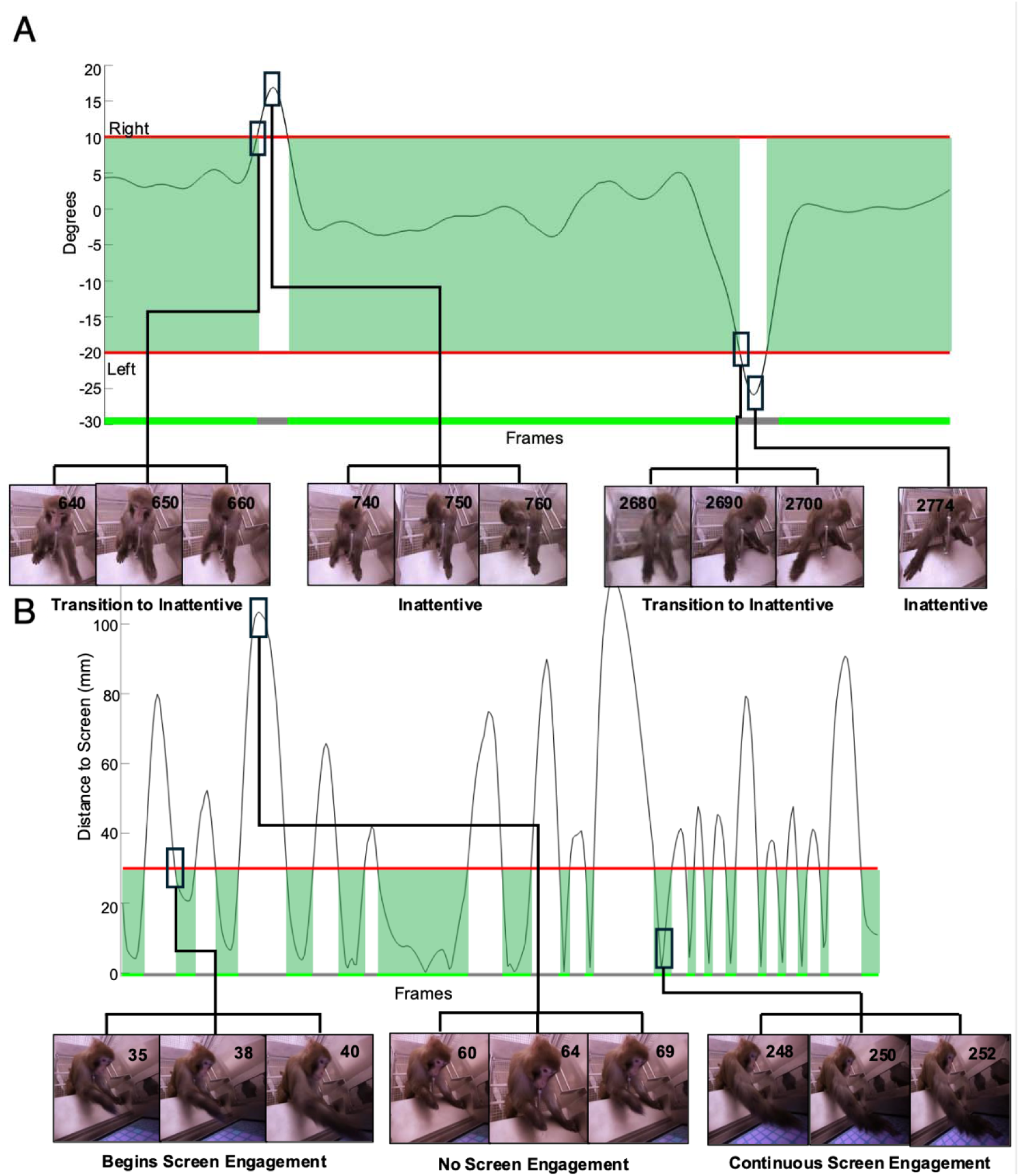
Example Classification of Attentiveness and Screen Engagement. (A) Analysis of NHP attentiveness over a 2-minute video segment using yaw angle as the primary feature. The black line represents the yaw angle, with red horizontal lines indicating left and right thresholds for attentiveness. Green bars denote attentive periods (yaw within thresholds), while grey bars indicate inattentive periods (yaw beyond thresholds). Image sequences at frames 640–660 and 2680–2700 show transitions from attentive to inattentive states, while frames 740-760 capture the NHP not being attentive. (B) Screen engagement analysis over a 30-second segment, based on the right wrist’s proximity to the touchscreen. The red horizontal line denotes the engagement threshold. Green bars highlight active engagement (distance below threshold), while distance above the threshold indicate disengagement. Frames 35–40 show the NHP initiating engagement, frames 60–69 depict disengagement, and frames 248–252 capture active screen engagement.

### 2.7. Classification of screen engagement

Screen engagement was assessed by analyzing the NHP’s head and arm movements in relation to the screen. Engagement is classified based on the calculated distance between the labeled joints of the arms and the hand relative to the screen. When the hand and arm movements cross a set threshold, the NHP is considered engaged with the screen. An example analysis is shown in **Figure 2B** over a 30-second video segment. The classification is based on the analysis of the NHP’s right wrist’s proximity to the touchscreen, represented by the black line tracing the distance of the wrist to the screen over time. The red horizontal line marks the touch threshold. When the black line falls below the red threshold line, the distance between the wrist and the screen indicates physical contact or near-touch interaction. These moments are classified as periods of active screen engagement and are highlighted by the green bars along the time axis. Conversely, when the black line remains above the red threshold, the NHP is considered disengaged from the screen, as its wrist is too far away to indicate an interactive gesture. The example frames illustrate specific moments that align with significant features of the distance trace (**Figure 2B**). For instance, frames 35, 38, and 40 show the NHP moving its hand toward the screen, corresponding to a decline in the black line towards the threshold, indicating the initiation of engagement. Frames 60, 64, and 69 capture the NHP moving his hand away from the screen, indicating a loss of engagement. In contrast, frames 248, 250, and 252 display the NHP’s hand engaging with the screen, indicating engagement with the screen.

### 2.8. Time course and task-specificity of attentiveness and screen engagement

The behavioral classification determines for each time frame of the video if the NHP is attentive or not and is engaged with the screen or not (**Figure 3B**). We analyzed this binary classification across time for different temporal window durations as well as across tasks that subjects performed during individual sessions (**Figure 3A**). The percentage of time being attentive and engaging with the screen can be directly translated into a score for the two metrics. For example, on a shorter time scale, attentiveness remained high over a ∼3 min epoch with only two smaller lapses (**Figure 3B**), while over a whole 90 min session, attentiveness scores systematically shifted varied, moving up when the first working memory task started, moving down when the effort control task started, and moving back up again when the second working memory task started (**Figure 3C**). This overall pattern was somewhat discernable but less apparent when we averaged the attentiveness scores for each task epoch across 31 experimental sessions **Figure 3D**. A more apparent task specificity was evident in the screen engagement scores (**Figure 3E**). Consistent with the higher rate of choices when performing the Maze-Learning task (each tile of a path had to be touched) and the effort control task (inflating the balloon required multiple touches per trial) the screen engagement was higher during these tasks compared to the working memory task (which required one choice per trial) (**Figure 3E**).

**Figure 3.**
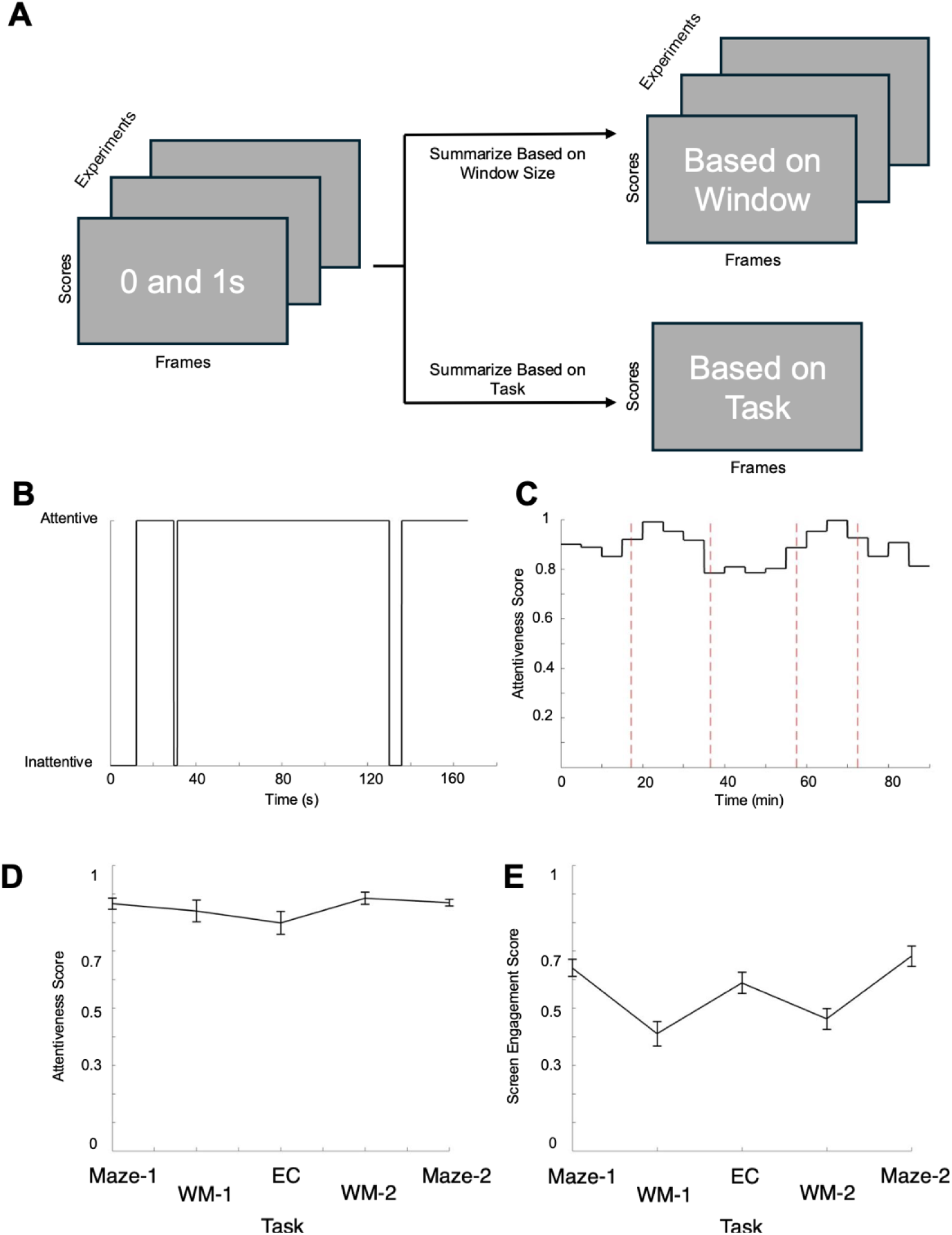
Visualization of Attentiveness and Screen Engagement Scores. (A) Schematic of the data processing pipeline for cross-trial analysis, summarizing attentiveness and screen engagement scores (0s and 1s) across frames, either by window size or by task (WM1, M1, EC, M2, WM2). (B) Example attentiveness score over a 5000-frame video segment, showing binary classification (attentive vs. inattentive). (C) Screen engagement score over a 90-minute session, averaged over 300 second windows, with red lines denoting transition between tasks. (D) Mean attentiveness scores per task (n = 31 sessions), with standard error bars representing variability across trials (E) Mean screen engagement scores per task (n = 31 sessions), with standard error bars representing variability across trials.

To quantify how well attentiveness and screen engagement scores distinguished NHP’s behavior during the performance of the different tasks we decoded the task labels using an unsupervised (k-means) clustering approach and a supervised (Random Forest) classifier (**Figure 4A**). K-means clustering is an unsupervised machine learning algorithm used to partition data into k distinct clusters based on feature similarity (Hartigan and Wong, 1979). It operates iteratively by assigning each data point to the nearest cluster centroid, recalculating centroids, and refining cluster assignments until convergence is reached. One of the key advantages of k-means is its efficiency in handling datasets with well-separated clusters, making it a popular choice for exploratory data analysis. The clustering was performed using Euclidean distance as the similarity metric. Random Forest is a supervised machine learning algorithm that operates as an ensemble of decision trees (Breiman, 2001). It works by constructing multiple decision trees during training and aggregating their predictions to improve accuracy and reduce overfitting. Each tree in the forest is trained on a random subset of the data using a technique known as bagging, and at each node, only a random subset of features is considered for splitting. This introduces diversity among trees, preventing over-reliance on specific patterns and enhancing the model’s generalizability. One advantage of Random Forest is its ability to handle overlapping and non-linearly separable data. Unlike the k-means clustering approach, which relies on predefined assumptions about the shape or density of data distributions, Random Forest builds multiple decision boundaries that adapt to complex feature interactions. When data points from different task categories exhibit significant overlap in attentiveness and engagement scores, individual decision trees may struggle to classify them correctly. However, by averaging predictions across many trees, Random Forest smooths out biases and effectively distinguishes patterns that may not be apparent in any single decision tree.

**Figure 4.**
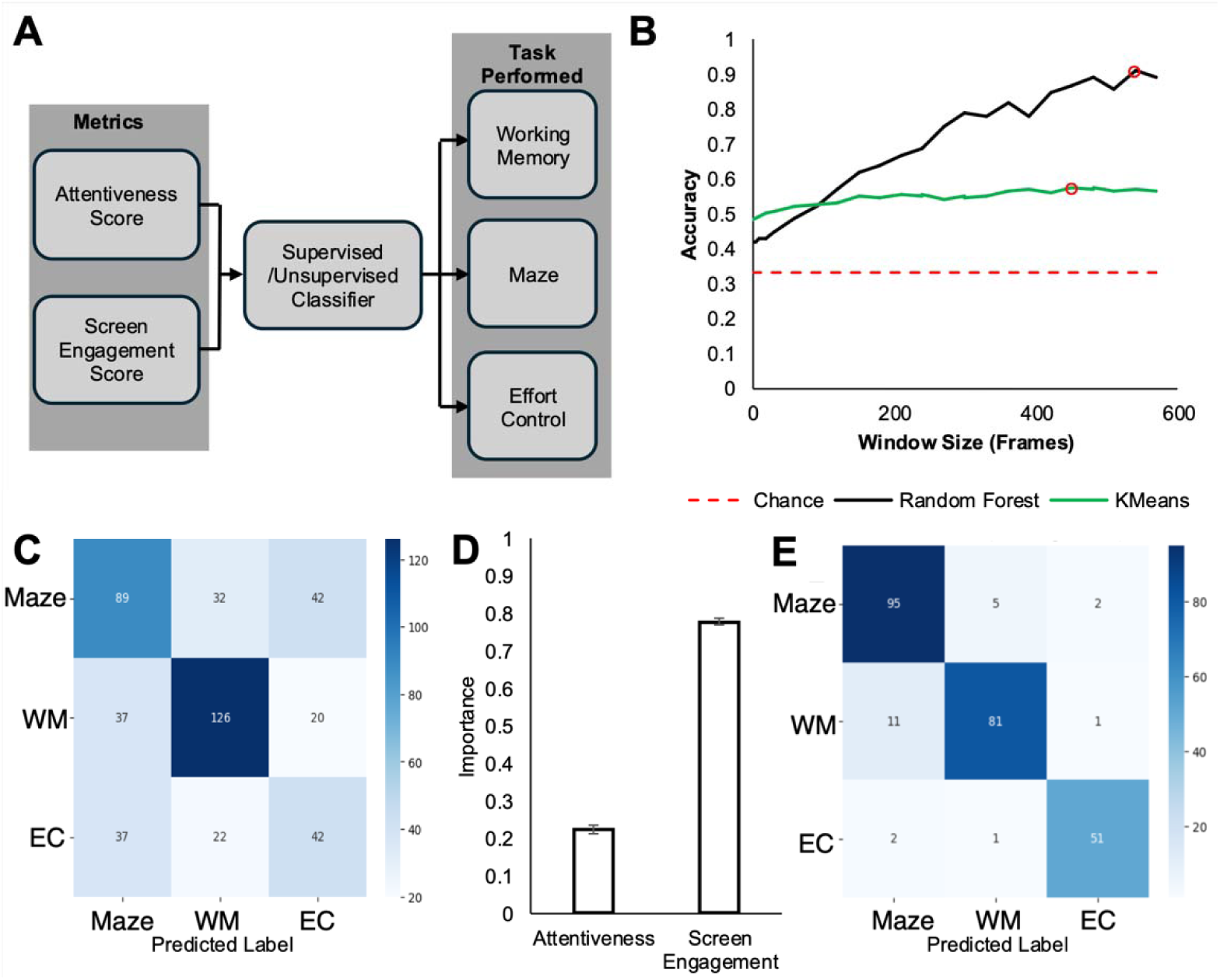
Task Classification Using Attentiveness and Screen Engagement Metrics. (A) Pipeline for task classification (Working Memory [WM], Maze [M], Effort Control [EC]) using attentiveness and screen engagement scores via supervised and unsupervised classifiers. (B) Classification accuracy versus window size, with Random Forest (black), K-Means (green), and chance (red dashed line); red circles mark peak accuracies. (C) Confusion matrix for K-Means at 450-frame window size, with accuracy of 0.575. (D) Feature importance for Random Forest at 540-frame window size, with attentiveness (0.222 ±0.007) and screen engagement (0.778 ±0.007). (E) Confusion matrix for Random Forest at 540-frame window size, with accuracy of 0.910.

We applied the classification to the reconstructed attentiveness and screen engagement time courses from 31 experimental sessions of one NHP. Before classification, the raw attentiveness and screen engagement data were preprocessed to ensure comparability across different sessions. Features were scaled and normalized using z-score normalization to account for inter-session variability. The dataset was parsed by the task type the NHP performed which were MZ, WM, and EC. To determine the optimal temporal window size to differentiate between task behaviors and evaluate the effect that window sizes have on classification accuracy we decoded the tasks using by averaging attentiveness and screen engagement scores at different temporal windows, ranging from single frames (30 ms) to 600 frames (20 sec) durations (**Figure 4B**). For the best decoding result, a feature importance analysis was used to evaluate the relative contribution of the screen engagement and attentiveness to the overall classification of the different tasks (**Figure 4D**).

## 3. Results

### 3.1. Attentiveness and screen engagement vary between cognitive tasks

Across thirty-one experimental sessions the attentiveness and screen engagement could be reliably extracted from video-captured performance of the cognitive tasks. Attentiveness scores remained high over extended periods of time (**Figure 3B**) and remained similarly high across tasks (one factorial ANOVA with the three tasks as factor, F= 1.45, p=0.221). In example sessions we observed that attentiveness scores remained high for ∼90% of the time during the initial maze task, increasing during the working memory task, dropping below 90% of the time during the effort control task, increasing to >90% during the second working memory task period, and leveling back to ∼90% using the second maze task at the end of the experimental session (**Figure 3C**). This temporal variation of attentiveness was evident across sessions, which quantifies that NHPs remained attentive towards the touchscreen throughout the sessions (**Figure 3D**).

A larger variability was seen for screen engagement, which quantified whether NHPs directed their hands and arms towards the screen. Screen engagement varied significantly across tasks (one factorial ANOVA, F= 9.98, p=3.476E-07) (**Figure 3E**). Screen engagement scores exceeded 50% of the total time in the maze tasks (proportion of time with high screen engagement in MZ1: 0.640, SE: 0.030; MZ2: 0.682, SE: 0.035), which involved touching continuously through the tiles of the path, and in the effort control task (EC: 0.588, SE: 0.038), which involved continuously touching the screen to inflate a virtual balloon in order to receive reward (**Figure 3E**). Screen engagement fell below 50% of the time during the working memory tasks (WM1: 0.411, SE: 0.043; WM2: 0.463, SE:0.037), which involved initiating individual trials and making one choice per trial after a variable short-term memory delay.

### 3.2 Classification of performing different cognitive tasks

We next used the attentiveness and screen engagement scores to quantify the temporal window that was needed to distinguish which cognitive task was performed (**Figure 4**). Decoding the tasks using k-means clustering showed a peak decoding accuracy of 57.5% (chance performance: 33%) at a time window size of 450 frames (15 s), with only subtle variations across 2-20 s time windows. Confusion matrices obtained at the time window with peak decoding performance (450 frames) showed that the WM task had the highest classification accuracy, the Maze-Learning task had the second highest and the Effort Control task the lowest classification accuracy (**Figure 4C**).

In contrast to moderate k-means clustering decoding of cognitive tasks, the Random Forrest classification steadily increased from ∼40-50% accuracy at high temporal resolution (e.g. 2 s, 60 frames) to above 90% decoding accuracy of the tasks when a time window of 18 s (540 frames) was used to average attentiveness and screen engagement scores (**Figure 4B**). A feature importance analysis at peak performance (91%, 540 frames) showed that the screen engagement scores were three times more important than the attentiveness scores for predicting which task was being performed (importance of 0.778 vs. 0.222, **Figure 4D**). Cross-validation by randomly selecting different percentages of the dataset was performed to validate the feature importance values (STD: 0.007). Analysis of the confusion matrix showed the highest accuracy for the Maze-Learning task, followed by the WM task and the Effort Control task (**Figure 4E**).

## 4. Discussion

Here, we introduced an integrated analysis and classification pipeline for time-resolved extraction of attentiveness and screen engagement scores of subjects performing different touchscreen based cognitive tasks. We found that that attentiveness and screen engagement scores reliably distinguished at >90% accuracy which of three tasks were performed. While tasks were classified above chance already at ∼ 2 sec temporal resolution, the maximal reliable classification required averaging attentiveness and screen engagement scores within 18 s (**Figure 4B**).

These findings establish a tool to empirically quantify cognitive variables from video by combining (1) state-of-the art pose estimation using DeepLabCut, (2) the 3D reconstruction of the task assessment environment, (3) a subject-specific thresholding of head/yaw orientation and arm/wrist/hand distance to the screen, and (4) time-resolved classification of attentiveness’ and screen engagement. The results shows that a 3D reconstruction of the head orientation and left and right yaw angle towards the screen serves as a robust signal to estimate subjects’ attention toward the screen. Attentiveness stayed at a high ∼90% throughout the duration of the task performance within the session and varied only moderately within a ∼5% range between tasks with low variability across sessions (**Figure 3D**). These characteristics suggest that attentiveness scores will be suited to distinguish subjects with different degrees of attentiveness’, the same subjects in different states, or subjects that are treated with pharmacological compounds that influence attentiveness (Millan et al., 2012; Azimi et al., 2020; Hassani et al., 2023). While our goal was to estimate attentiveness towards the screen, future studies could combine the head/yaw orientation information with gaze reconstruction to infer which objects on the screen were looked at. Recent progress in reconstructing gaze in monkeys suggest that such an approach will be viable with special tracking devices mounted on the subject (Singh et al., 2025).

Another insight from out study is that the combination of shoulder, arm, and wrist reconstruction provides a robust signal for evaluating the hand distance to the screen. We inferred screen engagement of the subject using this distance measure and showed that it is an informative marker for how often and how long subjects engaged with objects on the screen in tasks with varying requirement to engage with task elements like tile of a maze (Maze-Learning task), ballon outlines (Effort Control task), or individual objects (Working Memory task). We chose to estimate the wrist distance to the screen because it was less variable and less noisy than tracking knuckles of the individual digits and because it was more easily generalizable across subject whose specific touch patterns varying from using individual digits to multiple digits. The proposed quantification of screen engagement promises to be a versatile marker for the degree of cognitive engagement and cognitive processing speed, which are important variables varying with age and cognitive state, amongst others (Millan et al., 2012). One possible future extension towards developing behavioral markers of cognitive states is the integration of the video-based estimate of screen engagement with the behavioral accuracy and choice reaction times of the performed task. While our goal was to quantify how well tasks can be distinguished from video alone, future studies may be able to relate the vigor, precision, or consistency of screen engagement with the behavioral accuracy of performing different cognitive tasks, with the neuronal encoding of task relevant information, or with the specific effects of pharmacological intervention, electrical or optogenetic stimulation (Schweihoff et al., 2021). Such an integration of video-based behavioral analysis into a comprehensive neuro-behavioral assessment framework is a major goal of large-scale efforts to understand how brain circuits organize goal directed natural behavior (Voloh et al., 2023).

Taken together, our study introduced an analysis tool for assessing cognitive variables in NHPs from video alone. The adoption of real-time pose estimation tools like DeepLabCut in this study reflects recent advancements in behavioral neuroscience, which have enabled precise tracking of animal movements for cognitive analysis (Kane et al., 2020). By providing a quantitative and scalable method for behavioral assessment, the provided analysis framework opens new avenues for neurobehavioral research, facilitating a deeper understanding of cognition in NHPs across various experimental conditions.

## Appendix

The full codebase used for this study, including video preprocessing, 3D reconstruction, DeepLabCut pose estimation, behavioral classification, and data visualization, has been made publicly available to promote transparency and reproducibility. The repository includes MATLAB and Python scripts, documentation, and example datasets required to replicate the analysis pipeline described in this paper. The code for this project can be accessed at: https://github.com/ansencheung/Video-Analysis-and-Decoding-Pipeline.git

## Acknowledgments

This work was supported by the National Institute of Mental Health (R01MH129641 to TW). The funders had no role in study design, data collection and analysis, the decision to publish, or the preparation of this manuscript.

---

**Supplementary Figure S1.**
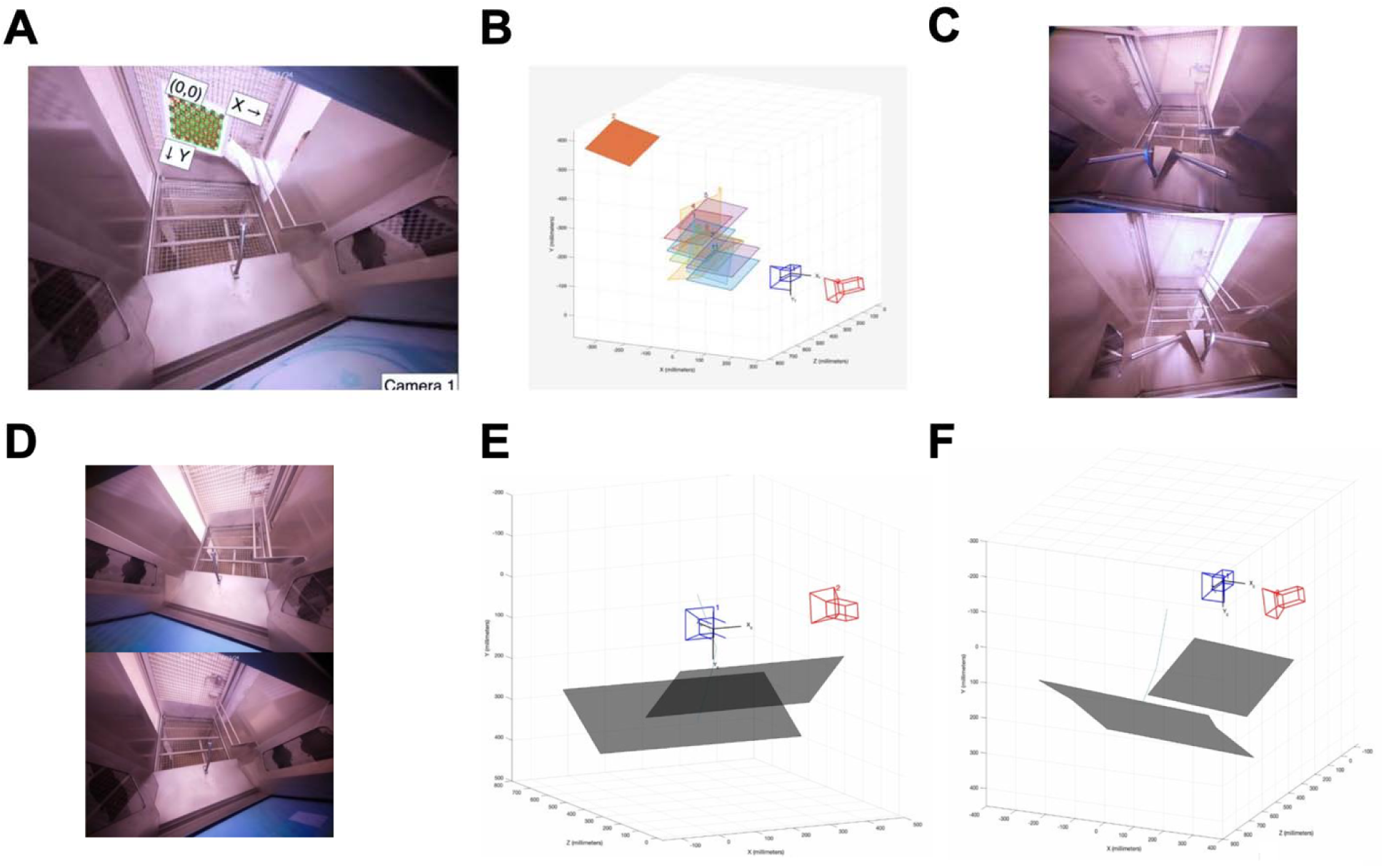
Static Kiosk Environment 3D Reconstruction. (A) Calibration of MATLAB Stereo Vision Toolbox with checkerboard. (B) Evaluation of calibration results of 11 checkerboard frames. (C) Left (bottom) and right (top) view of the old kiosk environment. (D) Left (bottom) and right (top) view of the new kiosk environment. (E) 3D reconstruction of the old kiosk environment with relative camera position, screen, base, and reward tube plotted. (F) 3D reconstruction of the new kiosk environment with relative camera position, screen, base, and reward tube plotted.

**Supplementary Figure S2.**
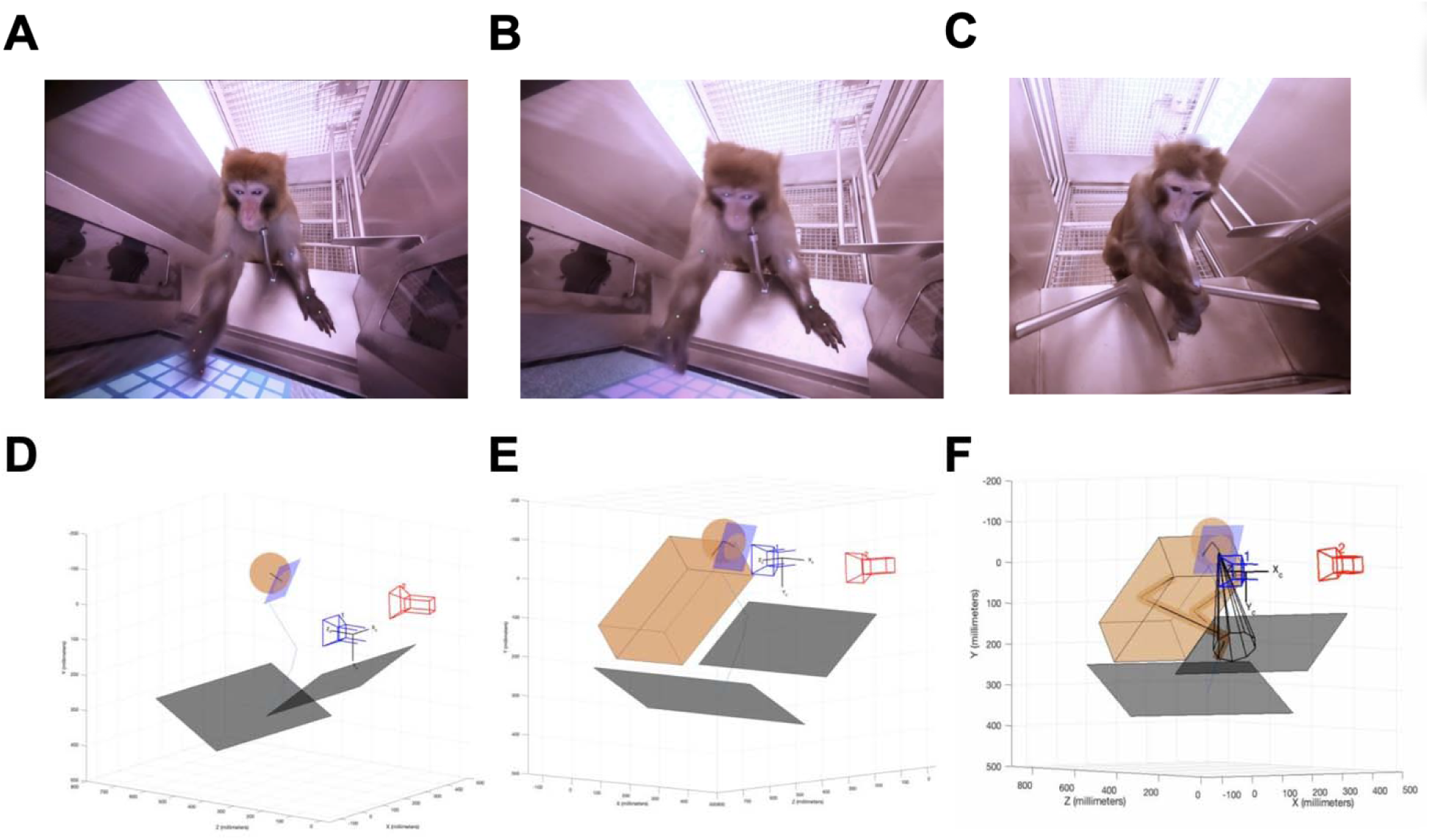
NHP 3D Reconstruction. (A) Manual labeled (11 body parts) of NHP (B) DLC Predicted (11 body parts) of NHP (C) Reference Frame for D, E, and F (D) 3D reconstruction of NHP head using predicted nose, left eye, and right eye 3D coordinates of NHP from reference frame. The three points create the blue plane. Head center and are estimated based on interpupillary distance. (E) 3D reconstruction of NHP torso based on head center location and screen position, assuming NHP is always facing screen. (F) 3D reconstruction of NHP arms and hands based on DLC predicted body parts.

## References

Azimi M, Oemisch M, Womelsdorf T (2020) Dissociation of nicotinic alpha7 and alpha4/beta2 sub-receptor agonists for enhancing learning and attentional filtering in nonhuman primates. Psychopharmacology (Berl) 237:997–1010.

Bain M, Nagrani A, Schofield D, Berdugo S, Bessa J, Owen J, Hockings KJ, Matsuzawa T, Hayashi M, Biro D, Carvalho S, Zisserman A (2021) Automated audiovisual behavior recognition in wild primates. Science Advances 7.

Bala PC, Eisenreich BR, Yoo SBM, Hayden BY, Park HS, Zimmermann J (2020) Automated markerless pose estimation in freely moving macaques with OpenMonkeyStudio. Nature Communications 11.

Biderman D et al. (2024) Lightning Pose: improved animal pose estimation via semi-supervised learning, Bayesian ensembling and cloud-native open-source tools. Nat Methods 21:1316–1328.

Breiman L (2001) Random Forests. Machine Learning 45:5–32.

Brookes O et al. (2024) PanAf20K: A Large Video Dataset for Wild Ape Detection and Behaviour Recognition. Int J Comput Vision 132:3086–3102.

Fuermaier ABM, Tucha L, Guo N, Mette C, Muller BW, Scherbaum N, Tucha O (2022) It Takes Time: Vigilance and Sustained Attention Assessment in Adults with ADHD. Int J Environ Res Public Health 19.

Grant JE, Chamberlain SR (2023) Attentional problems occur across multiple psychiatric disorders and are not specific for ADHD. CNS Spectr 28:357–360.

Hartigan JA, Wong MA (1979) A K-Means Clustering Algorithm. Journal of the Royal Statistical Society Series C: Applied Statistics 28:100–108.

Hassani SA, Lendor S, Neumann A, Sinha Roy K, Banaie Boroujeni K, Hoffman KL, Pawliszyn J, Womelsdorf T (2023) Dose-Dependent Dissociation of Pro-cognitive Effects of Donepezil on Attention and Cognitive Flexibility in Rhesus Monkeys. Biol Psychiatry Glob Open Sci 3:68–77.

Insafutdinov E, Pishchulin L, Andres B, Andriluka M, Schiele B (2016) DeeperCut: A Deeper, Stronger, and Faster Multi-person Pose Estimation Model. Lect Notes Comput Sc 9910:34–50.

Kane GA, Lopes G, Saunders JL, Mathis A, Mathis MW (2020) Real-time, low-latency closed-loop feedback using markerless posture tracking. Elife 9.

Kano F, Tomonaga M (2013) Head-mounted eye tracking of a chimpanzee under naturalistic conditions. PLoS One 8:e59785.

Keshavarzi S, Bracey EF, Faville RA, Campagner D, Tyson AL, Lenzi SC, Branco T, Margrie TW (2022) Multisensory coding of angular head velocity in the retrosplenial cortex. Neuron 110:532–543 e539.

Labuguen R, Matsumoto J, Negrete SB, Nishimaru H, Nishijo H, Takada M, Go Y, Inoue KI, Shibata T (2020) MacaquePose: A Novel “In the Wild” Macaque Monkey Pose Dataset for Markerless Motion Capture. Front Behav Neurosci 14:581154.

Li C, Xiao Z, Li Y, Chen Z, Ji X, Liu Y, Feng S, Zhang Z, Zhang K, Feng J, Robbins TW, Xiong S, Chen Y, Xiao X (2023) Deep learning-based activity recognition and fine motor identification using 2D skeletons of cynomolgus monkeys. Zool Res 44:967–980.

Liu Y, Wang M, Hou S, Wang X, Shi B (2025) Deep Learning-Based Markerless Hand Tracking for Freely Moving Non-Human Primates in Brain–Machine Interface Applications. Electronics 14.

Marshall JD, Li T, Wu JH, Dunn TW (2022) Leaving flatland: Advances in 3D behavioral measurement. Curr Opin Neurobiol 73:102522.

Marshall JD, Aldarondo DE, Dunn TW, Wang WL, Berman GJ, Olveczky BP (2021) Continuous Whole-Body 3D Kinematic Recordings across the Rodent Behavioral Repertoire. Neuron 109:420–437 e428.

Mathis A, Mamidanna P, Cury KM, Abe T, Murthy VN, Mathis MW, Bethge M (2018) DeepLabCut: markerless pose estimation of user-defined body parts with deep learning. Nat Neurosci 21:1281–1289.

Millan MJ et al. (2012) Cognitive dysfunction in psychiatric disorders: characteristics, causes and the quest for improved therapy. Nat Rev Drug Discov 11:141–168.

North R, Wurr R, Macon R, Mannion C, Hyde J, Torres-Espin A, Rosenzweig ES, Ferguson AR, Tuszynski MH, Beattie MS, Bresnahan JC (2021) Quantifying the kinematic features of dexterous finger movements in nonhuman primates with markerless tracking. 43rd Annual International Conference of the IEEE Engineering in Medicine & Biology Society (EMBC) 6110–6115.

Ottenheimer DJ, Bari BA, Sutlief E, Fraser KM, Kim TH, Richard JM, Cohen JY, Janak PH (2020) A quantitative reward prediction error signal in the ventral pallidum. Nat Neurosci 23:1267–1276.

Pereira TD et al. (2022) SLEAP: A deep learning system for multi-animal pose tracking. Nature Methods 19:486–+.

Redish AD (2016) Vicarious trial and error. Nat Rev Neurosci 17:147–159.

Ryan AM, Freeman SM, Murai T, Lau AR, Palumbo MC, Hogrefe CE, Bales KL, Bauman MD (2019) Non-invasive Eye Tracking Methods for New World and Old World Monkeys. Front Behav Neurosci 13:39.

Schweihoff JF, Loshakov M, Pavlova I, Kuck L, Ewell LA, Schwarz MK (2021) DeepLabStream enables closed-loop behavioral experiments using deep learning-based markerless, real-time posture detection. Commun Biol 4:130.

Singh VP, Li J, Dawson K, Mitchell JF, Miller CT (2025) Active vision in freely moving marmosets using head-mounted eye tracking. Proc Natl Acad Sci U S A 122:e2412954122.

Vogg R, Luddecke T, Henrich J, Dey S, Nuske M, Hassler V, Murphy D, Fischer J, Ostner J, Schulke O, Kappeler PM, Fichtel C, Gail A, Treue S, Scherberger H, Worgotter F, Ecker AS (2025) Computer vision for primate behavior analysis in the wild. Nat Methods.

Voloh B, Maisson DJN, Cervera RL, Conover I, Zambre M, Hayden B, Zimmermann J (2023) Hierarchical action encoding in prefrontal cortex of freely moving macaques. Cell Reports 42.

Watson MR, Voloh B, Naghizadeh M, Womelsdorf T (2019) Quaddles: A multidimensional 3-D object set with parametrically controlled and customizable features. Behav Res Methods 51:2522–2532.

Watson MR, Traczewski N, Dunghana S, Boroujeni KB, Neumann A, Wen X, Womelsdorf T (2023) A Multi-task Platform for Profiling Cognitive and Motivational Constructs in Humans and Nonhuman Primates. bioRxiv.

Wei SE, Ramakrishna V, Kanade T, Sheikh Y (2016) Convolutional Pose Machines. Proc Cvpr Ieee:4724–4732.

Wen X, Neumann A, Dhungana S, Womelsdorf T (2025) Flexible Learning and Re-ordering of Context-dependent Object Sequences in Non-human Primates. bioRxiv 10.1101/2024.11.24.625056.

Wiltschko AB, Tsukahara T, Zeine A, Anyoha R, Gillis WF, Markowitz JE, Peterson RE, Katon J, Johnson MJ, Datta SR (2020) Revealing the structure of pharmacobehavioral space through motion sequencing. Nat Neurosci 23:1433–1443.

Womelsdorf T, Thomas C, Neumann A, Watson MR, Banaie Boroujeni K, Hassani SA, Parker J, Hoffman KL (2021) A Kiosk Station for the Assessment of Multiple Cognitive Domains and Cognitive Enrichment of Monkeys. Front Behav Neurosci 15:721069.

Xing F, Sheffield AG, Jadi MP, Chang SWC, Nandy AS (2024) Automated 3D analysis of social head-gaze behaviors in freely moving marmosets. BioRxiv.

